# Phosphate sensing by PhoPR regulates the cytotoxicity of *Staphylococcus aureus*

**DOI:** 10.1101/2025.06.24.661297

**Authors:** Nathanael Palk, Tarcisio Brignoli, Marcia Boura, Ruth C. Massey

## Abstract

*Staphylococcus aureus* has evolved a complex regulatory network to coordinate expression of virulence factors, including cytolytic toxins, with host environmental signals. Central to this network are two-component systems, in which a histidine kinase senses an external signal and activates a response regulator via phosphorylation, leading to changes in gene expression. Using a comprehensive screen of transposon mutants in each of the non-essential histidine kinase and response regulatory genes in *S. aureus*, we demonstrate that 11 of these 16 systems regulate cytotoxicity. Further characterisation of a *phoP* mutant revealed that its impact on cytotoxicity is mediated through the Agr quorum-sensing system. Notably, we found that unphosphorylated PhoP is an activator of Agr activity, while phosphorylated PhoP also acts as a weak activator of Agr activity in high phosphate environments but as a repressor in low phosphate environments. Overall, we have demonstrated that phosphate sensing through PhoPR is a novel regulator of cytotoxicity in *S. aureus*. Moreover, our study challenges the canonical model of TCSs as simple on/off systems and highlights the importance of unphosphorylated response regulators in gene regulation.

**Importance:** The production of cytolytic toxins is the major means by which bacterial pathogens damage host tissue and cause disease. Understanding the activity and regulation of these toxins is critical for the identification of means to block them and prevent the development of disease. In this study we focused on a specific regulatory mechanism, the two-component systems (TCSs), that enable bacteria to sense their environment and adapt accordingly. In the traditional model of a TCS, a response regulator (RR) is phosphorylated by a histidine kinase (HK), which enables it to activate or repress expression of target genes, which may include toxins or regulators of toxins. We found that 11 of *S. aureus’* 16 TCSs affect toxin production, highlighting that *S. aureus* integrates a broad range of environmental cues to regulate toxicity. We focused on one of these TCSs, the PhoPR system and found that sensing of inorganic phosphate is a novel regulator of cytotoxicity in *S. aureus*. Furthermore, we found that the RR of this system acts as a strong activator of toxicity in its unphosphorylated form, challenging the traditional model of a TCS as only active upon signal activation.

## Introduction

*Staphylococcus aureus* is a Gram-positive bacterium and opportunistic human pathogen which was associated with more than one million deaths globally in 2019 (1). It is a leading cause of a wide range of infections, including skin and soft tissue infections, bacteraemia, endocarditis, pneumonia and osteomyelitis (2). The ability of *S. aureus* to cause a diverse spectrum of diseases is generally attributed to an impressive arsenal of virulence factors (3-6). These include several cytolytic toxins, which lyse host cells by forming pores or channels in their membranes, referred to from herein as cytotoxicity. Alpha-haemolysin (Hla), the bicomponent leukotoxins (LukSF-PV, LukAB, and LukED), and gamma haemolysin (HlgAB and HlgCB) use a receptor-dependent mechanism to cause lysis. In contrast, phenol-soluble modulins (PSMs) have detergent-like properties (7-9). These have been linked to several aspects of *S. aureus* pathogenesis, including immune evasion, nutrient acquisition and inflammation (8, 10, 11). Importantly, cytolytic toxins are expressed in coordination with host environmental signals through a complex regulatory network and understanding this is key to linking their pathogenic role to specific disease states (12-14).

In bacteria, two component systems (TCSs) are a widespread mechanism used to sense and respond to environmental signals (15). In response to a specific signal, a membrane-bound histidine kinase (HK) will undergo ATP-dependent autophosphorylation. Subsequently, the phosphoryl group will be transferred to a cytoplasmic DNA-binding response regulator (RR) (16). Phosphorylation of the RR induces a conformation change and modulates its affinity for gene promoters, where they can act as transcriptional activators or repressors (17). Some TCSs also contain auxiliary proteins which can be involved in sensing or signal transduction (18, 19).

The genome of *S. aureus* contains 16 TCSs, with several of these linked to regulating expression of cytolytic toxins. For example, the accessory gene regulator (Agr) TCS uses quorum sensing to upregulate most *S. aureus* toxins, including Hla, leukotoxins, delta-hemolysin (Hld) and PSMs (20-22). The staphylococcal accessory element (Sae) TCS responds to phagocytosis-related signals to upregulate expression of Hla, leukotoxins and HlgAB and HlgCB (23-26). Similarly, the autolysis-related locus (Arl) TCS acts as a metabolic sensor and induces expression of leukotoxins in response to nutritional immunity (13, 27-29). The remaining TCSs have been linked to cell wall biosynthesis and antibiotic resistance (VraRS, GraRS, NsaRS), autolysis (WalRK, LytRS), respiration (SrrBA, AirRS, and NreCB), and nutrient sensing (HssRS, KdpDE, and PhoPR) (17). Additionally, there is a TCS with a currently unknown function (DesRK).

Current knowledge of the TCSs which regulate cytotoxicity in *S. aureus* is mainly derived from transcriptional comparisons between wild-type strains and mutant pairs in histidine kinase (HK) and/or response regulator (RR) genes. However, we are currently lacking a comprehensive study which evaluates and compares the contribution of all the TCSs in *S. aureus* to a single phenotype such as cytotoxicity. To address this, we screened the cytotoxicity of transposon mutants in each non-essential HK and RR gene in the *S. aureus* NTML reference CA-MRSA strain JE2. Using this approach, we identified several TCS mutants with reduced cytotoxicity, including a mutant of the *phoP* gene, the RR from the PhoPR TCS. The *phoP* mutant has a lower abundance of PSMs in the bacterial supernatant due to decreased Agr activity. Interestingly, we found that PhoP regulation of Agr activity is differential depending on the concentration of inorganic phosphate (Pi) in growth media. Furthermore, we show that the phosphorylation state of PhoP is a key factor in mediating this differential regulation.

## Methods

### Bacterial Strains and Growth Conditions

A list of the *S. aureus* strains, plasmids and primers used in this study are available in Tables 1, 2 and 3. *S. aureus* strains were grown at 37°C on tryptic soy agar (TSA) or in tryptic soy broth (TSB) with shaking at 180rpm. Mutant strains of all HK and RR genes in the genome of USA300 JE2 were obtained from the Nebraska Transposon Mutant Library (NTML) (30) and grown in media supplemented with erythromycin (5 μg/ml). *S. aureus* strains carrying the pRMC2 plasmid were selected with chloramphenicol (10 µg/ml) and expression of the inserted gene induced with anhydrotetracycline (aTC) (200 ng/ml).

**Table 1.** Strains used in this study.

| Strain | Description | Reference |
| --- | --- | --- |
| JE2 | USA300; CA -MRSA, type IV SCCmec; lacking plasmids p01 and p03; wild -type strain of the NTML | (30) |
| *agrA::tn* | *agrA* transposon mutant in JE2 | (30) |
| *agrC::tn* | *agrC* transposon mutant in JE2 | (30) |
| *saeR::tn* | *saeR* transposon mutant in JE2 | (30) |
| *saeS::tn* | *saeS* transposon mutant in JE2 | (30) |
| *arlR::tn* | *arlR* transposon mutant in JE2 | (30) |
| *arlS::tn* | *arlS* transposon mutant in JE2 | (30) |
| *graR::tn* | *graR* transposon mutant in JE2 | (30) |
| *graS::tn* | *graS* transposon mutant in JE2 | (30) |
| *vraR::tn* | *vraR* transposon mutant in JE2 | (30) |
| *vraS::tn* | *vraS* transposon mutant in JE2 | (30) |
| *nsaR::tn* | *nsaR* transposon mutant in JE2 | (30) |
| *nsaS::tn* | *nsaS* transposon mutant in JE2 | (30) |
| *lytR::tn* | *lytR* transposon mutant in JE2 | (30) |
| *lytS::tn* | *lytS* transposon mutant in JE2 | (30) |
| *kdpE::tn* | *kdpE* transposon mutant in JE2 | (30) |
| *kdpD::tn* | *kdpD* transposon mutant in JE2 | (30) |
| *phoP::tn* | *phoP* transposon mutant in JE2 | (30) |
| *phoR::tn* | *phoR* transposon mutant in JE2 | (30) |
| *hptR::tn* | *hptR* transposon mutant in JE2 | (30) |
| *hptS::tn* | *hptS* transposon mutant in JE2 | (30) |
| *hssR::tn* | *hssR* transposon mutant in JE2 | (30) |
| *hssS::tn* | *hssS* transposon mutant in JE2 | (30) |
| *srrA::tn* | *srrA* transposon mutant in JE2 | (30) |
| *srrB::tn* | *srrB* transposon mutant in JE2 | (30) |
| *nreC::tn* | *nreC* transposon mutant in JE2 | (30) |
| *nreB::tn* | *nreB* transposon mutant in JE2 | (30) |
| *airS::tn* | *airS* transposon mutant in JE2 | (30) |
| *desR::tn* | *desR* transposon mutant in JE2 | (30) |
| *desK::tn* | *desK* transposon mutant in JE2 | (30) |
| *phoP::tn* pRMC2 | *phoP* transposon mutant in JE2 transformed with empty pRMC2 vector | This study |
| *phoP::tn pphoP* | *phoP* transposon mutant complemented with *phoP* gene cloned into the pRMC2 expression plasmid | This study |
| JE2<br>pRNAIII::gfp | JE2 transformed with GFP reporter plasmid of RNAIII expression | This study |
| *phoP::tn*<br>*pRNAIII::gfp* | *phoP* transposon mutant transformed with GFP reporter plasmid of RNAIII expression | This study |
| *agrA::tn*<br>*pRNAIII::gfp* | *agrA* transposon mutant transformed with GFP reporter plasmid of RNAIII expression | This study |
| JE2 pAH5E | JE2 transformed with YFP reporter vector with no promoter driving *yfp* expression | This study |
| JE2 *ppstS::yfp* | JE2 transformed with YFP reporter plasmid of *pstS* expression | This study |
| *phoP::tn* pAH5E | *phoP* transposon mutant transformed with YFP reporter vector with no promoter driving *yfp* expression | This study |
| *phoP::tn*<br>*ppstS::yfp* | *phoP* transposon mutant transformed with YFP reporter plasmid of *pstS* expression | This study |
| *phoP::tn pphoP*<br>(D53A) | *phoP* transposon mutant complemented with *phoP* (D53A mutation) cloned into the pRMC2 expression plasmid | This study |
| *phoP::tn pphoP*<br>(D53E) | *phoP* transposon mutant complemented with *phoP* (D53E mutation) cloned into the pRMC2 expression plasmid | This study |

**Table 2.**
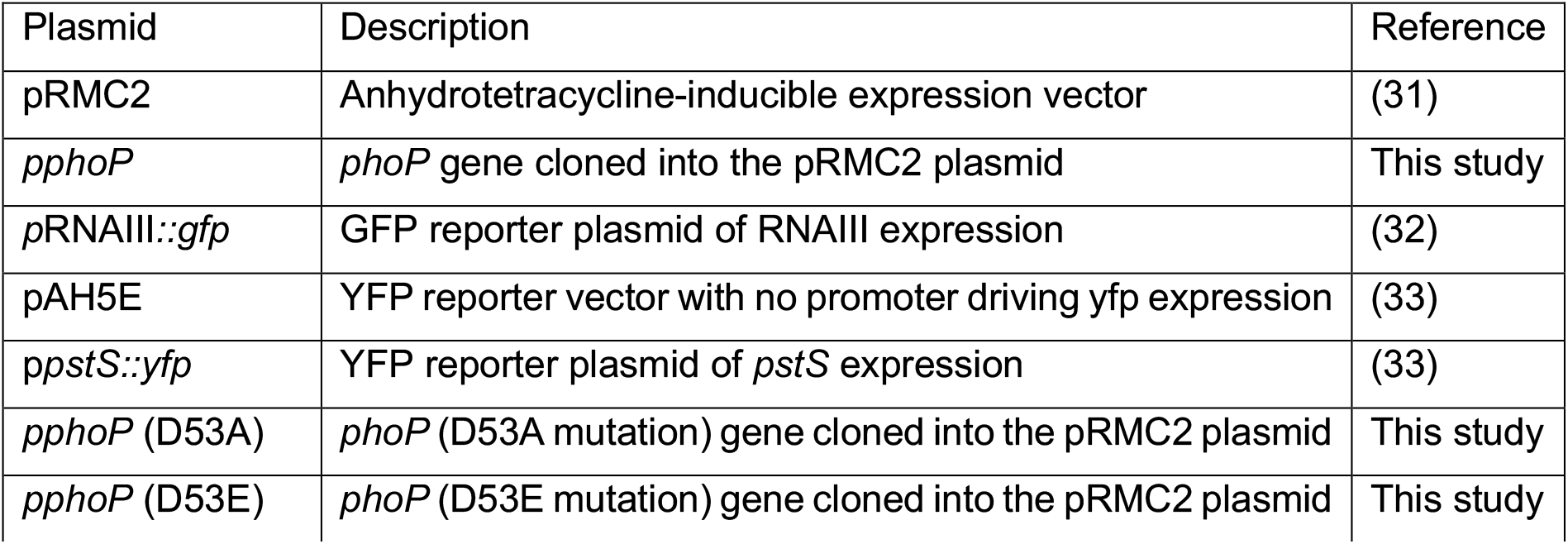
Plasmids used in this study.

| Plasmid | Description | Reference |
| --- | --- | --- |
| pRMC2 | Anhydrotetracycline-inducible expression vector | (31) |
| *pphoP* | *phoP* gene cloned into the pRMC2 plasmid | This study |
| *pRNAIII::gfp* | GFP reporter plasmid of RNAIII expression | (32) |
| pAH5E | YFP reporter vector with no promoter driving *yfp* expression | (33) |
| *ppstS::yfp* | YFP reporter plasmid of *pstS* expression | (33) |
| *pphoP* (D53A) | *phoP* (D53A mutation) gene cloned into the pRMC2 plasmid | This study |
| *pphoP* (D53E) | *phoP* (D53E mutation) gene cloned into the pRMC2 plasmid | This study |

**Table 3.**
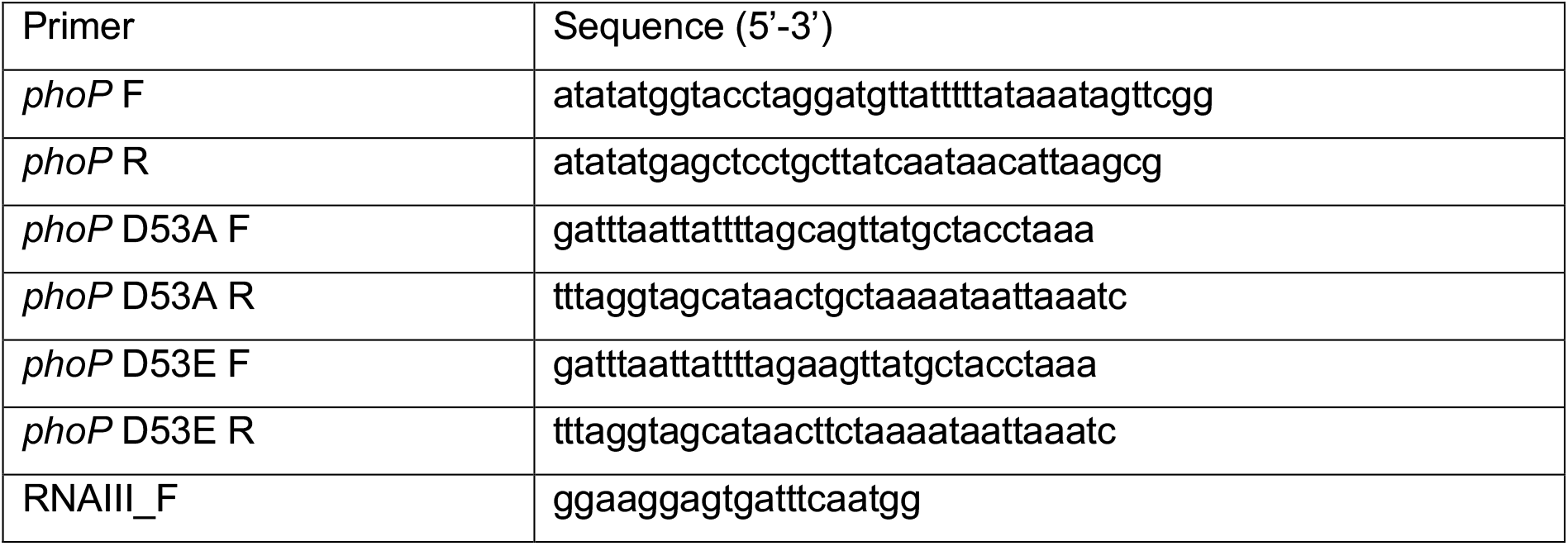
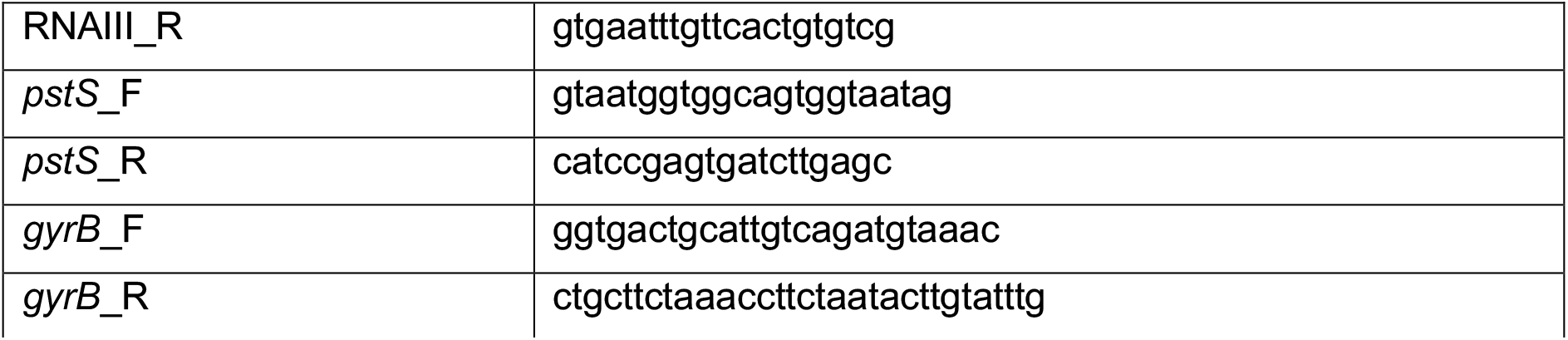
Primers used in this study.

### THP-1 Cytotoxicity Assay

THP-1 cells were exposed to the supernatant from *S. aureus* overnight cultures as a measure of cytotoxicity as they are sensitive to lysis by 13 of the 15 toxins present in the supernatant (34). THP-1 cells were sub-cultured every 2-3 days in RPMI 1640 supplemented with heat-inactivated fetal bovine serum (FBS) (10%), L-glutamine (1 μM), penicillin (200 U/ml), and streptomycin (0.1 mg/ml) at 37°C with 5% CO2. For the toxicity assay, cells were harvested by centrifugation (1250rpm, 20°C, 5 minutes) and resuspended in Hank’s Balanced Salt Solution (HBSS) (Biowhittaker) to a final density of 1 × 10^6^ to 1.5 × 10^6^ cells/ml. Bacterial supernatants were extracted after 18 hours of growth by centrifugation (4000rpm, 4°C, 5 minutes) and incubated with harvested THP-1 cells for 12 minutes at 37°C. Cell death was quantified via trypan blue exclusion with percentage (%) THP-1 killing defined as the number of dead THP-1 cells out of the total number of THP-1 cells counted for each sample. Each assay was performed with three technical replicates and three biological replicates. For comparisons across multiple assays, this percentage was taken relative to the THP-1 killing of the wild-type strain (JE2) in each assay to account for any variability in the sensitivity of the THP-1 cells. Incubation of THP-1 cells with TSB was used as a negative control to ensure appropriate viability of the THP-1 cells for each assay.

### Complementation of the *phoP::tn* mutant

To verify the *phoP::tn* mutation was responsible for the reduced cytotoxicity phenotype, the wild-type *phoP* gene was reintroduced into the transposon mutant via the pRMC2 vector. This vector was selected as it controls of gene expression through an aTC-inducible promoter.

A wild-type copy of the *phoP* gene was PCR amplified from JE2 genomic DNA with KpnI and SacI restriction sites using *phoP* F and *phoP* R primers (Table 3). The PCR product was checked for a product of the correct size by agarose gel electrophoresis and purified using the QIAquick® PCR purification kit (Qiagen). Purified *phoP* and pRMC2 vector were then digested with KpnI and SacI (NEB) restriction enzymes and the products purified again. Insert and plasmid DNA were ligated in a 3:1 molar ratio using T4 DNA ligase (NEB) to generate p*phoP*. This was transformed into *E. coli* Mach 1 competent cells via heat shock and successful transformants confirmed by colony PCR. Plasmids were isolated from Mach 1, passaged through *S. aureus* RN4220 and then transformed into *phoP::tn* via electroporation. For electroporation, competent cells were generated by culturing strains to an OD_600nm_ of 0.5 and washing three times in 0.5M sucrose. For transformations, competent cells were incubated in ice with 100ng of plasmid DNA for 30 mins. Cells were transferred into 0.2mm electroporation curvettes (Bio-Rad) for electroporation (25 μF, 2.5 KV and 100 Ω; pulse time of 2.5 msec). Following electroporation, 700μL of TSB was added and cells were incubated at 37°C for 1 hour. Successful transformants were selected for by plating onto TSA with 10μg/ml chloramphenicol. To test for complementation of cytotoxicity, strains carrying the pRMC2 vector were cultured in TSB with 10μg/ml chloramphenicol and 200ng/ml anhydrotetracycline and supernatants extracted after 18 hours of growth. Cytotoxicity in *phoP::tn* p*phoP* was compared to a strain carrying the empty vector (*phoP::tn* pRMC2) as these could be grown under the same conditions.

### PSM Abundance

To extract the supernatant, 1ml of *S. aureus* overnight culture was centrifuged (13000rpm, 5 mins). The supernatant was then diluted 2-fold in TSB and combined with 5X protein loading dye (National Diagnostics). Samples were boiled at 98°C for 5 minutes and loaded on to 4-12% nUView Tris-Glycine Precast Gels. Protein was separated at 100-130V until the dye front had run off the gel. Protein bands were visualised using Quick Coomassie Stain (Protein Ark) overnight and gels destained in deionised water for at least 5 hours. Band intensities were quantified using ImageJ.

### GFP/YFP reporter assays

Reporter plasmids (pRNAIII::*gfp*, pAH5E or p*pstS::yfp*) were transformed into JE2, *phoP::tn* or *agrA::tn* via electroporation. To determine RNAIII expression in TSB, strains with pRNAIII::*gfp* were grown overnight in TSB with 10ug/ml chloramphenicol and normalised to an OD_600nm_ of 0.05 in fresh TSB. GFP fluorescence (485nm excitation/520nm emission/1000 gain) and OD_600nm_ readings were taken every 30 mins for a 24-hour period with 200rpm shaking in between readings. Measurements were taken in a black 96 well plate (Costar) using a CLARIOstar plate reader. GFP fluorescence (485nm excitation/520nm emission/1500 gain) and these were normalised to OD_600nm_ readings (F/OD).

To determine RNAIII and *pstS* expression under different phosphate concentrations, overnight cultures of strains carrying either *p*RNAIII*::gfp* or *ppstS::yfp* were washed and resuspended in phosphate-depleted RPMI media (MP Biomedicals). Strains were then normalised to an OD_600nm_ of 0.05 in phosphate-depleted RPMI media with phosphate levels adjusted using 0.2M sodium phosphate buffer, pH 7.4 (Thermofisher) and incubated at 37°C overnight with shaking. A 200ul volume of culture was aliquoted into a black 96 well plate (Costar). YFP measurements (485nm excitation/520nm emission/1500 gain) were normalised to OD_600nm_ and the F/OD of a corresponding strain carrying empty pAH5E was subtracted from this value as described previously (33).

### Site-directed Mutagenesis

Site-directed mutagenesis was performed to mutate wild-type *phoP* in p*phoP* to two variants: D53A and D53E. PCR amplification was performed using p*phoP* from Mach1 *E. coli* as template DNA with *phoP* D53A F and *phoP* D53A R or *phoP* D53E F and *phoP* D53E R primers (Table 3). The PCR product was checked for a band of the correct size by agarose gel electrophoresis. 1ul of DpnI was then added and the PCR product was incubated at 37°C for 1 hour to remove template DNA. 5ul of the product was then transformed into Mach 1, passaged through *S. aureus* RN4220 and then transformed into *phoP::tn* via electroporation.

### mRNA Extraction and qRT-PCR

*S. aureus* strains were grown to an OD_600nm_ of 4 in 50ml TSB and incubated with 200ng/ml aTC for 1 hour at 37°C with shaking at 180 rpm. 2ml of culture was then collected and RNA stabilised using RNAprotect bacterial reagent (Qiagen). RNA extractions were performed as described previously (35) using the Quick-RNA fungal/bacterial miniprep kit (Zymo research). Genomic DNA was removed from RNA samples using a TURBO DNA-free kit (Thermo Fisher), and reverse transcription was performed using the qScript cDNA synthesis kit (Quantabio). To verify samples were free from DNA contamination, a reverse transcription negative reaction was performed on all RNA samples and qRT-PCR performed. Threshold values were then compared to reverse transcription positive samples. Real-time PCR was performed using a KAPA SYBR fast qPCR kit (Kapa Biosystems), using RNAIII_F and RNAIII_R, *pstS*_F and *pstS*_R or *gyrB*_F and *gyrB*_R. *gyrB* was used as the housekeeping gene. All primer sets were validated using 5-fold dilutions of JE2 genomic DNA with efficiencies of 104.19, 106.19%, and 100.67% for *pstS*, RNAIII and *gyrB*, respectively.

### Statistical Analysis

Statistical comparisons between two samples were performed with an unpaired two-tailed t-test using GraphPad Prism. A *p* value of < 0.05 was considered significant. In our screening of cytotoxicity, multiple *t*-tests were used and a false discovery rate (FDR) of 1% was applied account for multiple comparisons. Therefore, samples were considered significant if they had an adjusted *p*-value > 0.01. Data is displayed as the mean ± standard deviation of the mean (SD) and experiments were performed with three biological replicates.

## Results

### Cytotoxicity is altered in transposon mutants of many TCSs in *S. aureus*

To identify the TCSs in *S. aureus* that regulate cytotoxicity, we used transposon insertion mutants in the histidine kinase (HK) and response regulator (RR) genes in the genome of *S. aureus* JE2 from the Nebraska Transposon Mutant Library (NTML) (30). As the WalKR TCS is essential for cell viability, there was no mutant strain of the *walK* and *walR* genes available. Additionally, there was no mutant strain of the *airR* gene in the NTML. From a screening of 29 transposon mutants, 17 demonstrated a statistically significant difference in cytotoxicity compared to the wild-type strain (Fig. 1). All unadjusted and adjusted *p*-values are provided in Supplementary Table 1. Importantly, all transposon mutants in TCSs which have been demonstrated in previous studies to directly regulate expression of cytolytic toxins (*agrA::tn, agrC::tn, saeR::tn, saeS::tn, arlR::tn* and *arlS::tn*) had a significant reduction in THP-1 killing, providing proof of principle of our cytotoxicity assay (3, 12, 28). In addition to the established cytotoxicity linked TCSs, others involved in cell wall biosynthesis and autolysis (*graR::tn, graS::tn, nsaR::tn, nsaS::tn* and *lytR::tn*) had reduced cytotoxicity in our assay, indicating that there could be a link between regulation of the cell wall and cytotoxicity. Nutrient sensing also appeared to have an impact on cytotoxicity as several TCS mutants (*kdpE::tn, phoP::tn* and *hptS::tn*) had reduced cytotoxicity compared to the wild-type strain. In contrast, TCSs involved in respiration had a less of an impact on cytotoxicity, with only the *nreC::tn* mutant displaying reduced THP-1 killing. Interestingly, the HK and RR mutants of DesRK, a TCS with uncharacterised function, had reduced cytotoxicity compared to the wild-type strain. Overall, this data demonstrates that the majority of the TCSs in *S. aureus* contribute to regulating cytotoxicity.

**Figure 1.**
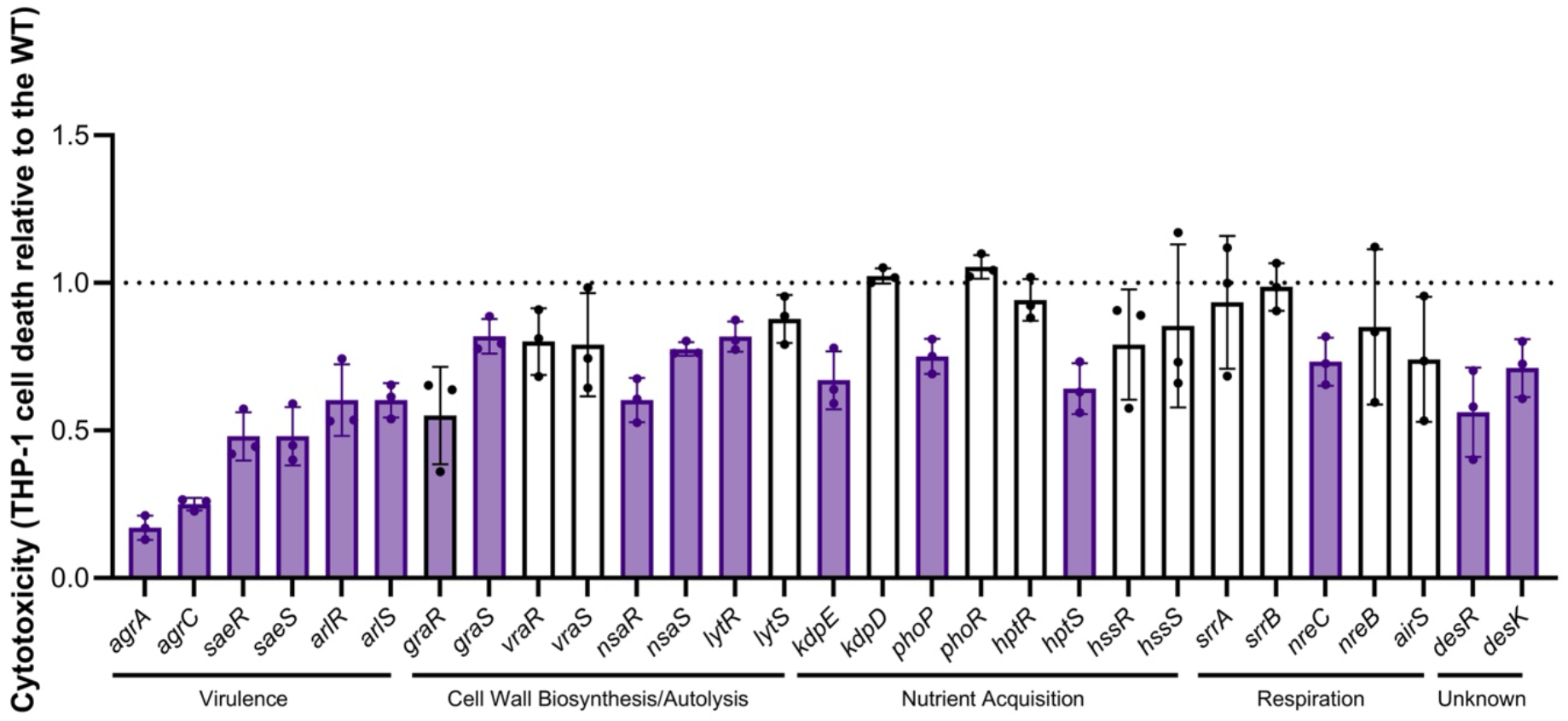
Cytotoxicity is altered in transposon mutants of TCSs in *S. aureus*. To quantify cytotoxicity, the bacterial supernatant from transposon mutants in each non-essential histidine kinase (HK) and response regulator (RR) gene was incubated with THP-1 cells and percentage of dead cells quantified using trypan blue exclusion. THP-1 killing was calculated relative to the THP-1 killing of JE2 (WT) in each assay. Each dot represents one biological replicate (n = 3), error bars the standard deviation, and statistical significance was determined using multiple *t*-tests with a false discovery rate (FDR) of 1% applied. Significant hits, which had an adjusted *p*-value ≤ 0.01 are highlighted in purple.

### The *phoP::tn* Mutant Produces a Lower Abundance of Phenol-Soluble Modulins

PhoPR is an inorganic phosphate (Pi) responsive TCS which is present in many bacterial species (36). PhoR, the HK of the system, undergoes autophosphorylation in response to Pi limitation and then phosphorylates PhoP, the RR, which can activate or repress target genes (36, 37). Whilst most of these genes are involved in phosphate homeostasis, the PhoPR TCS has been linked to regulating virulence factors in several other bacterial species such as *Escherichia coli* and *Mycobacterium tuberculosis* (38-40). Therefore, we wanted to establish whether phosphate sensing regulates cytotoxicity in *S. aureus*. Firstly, we sought to rule out any polar effects of the transposon insertion or spurious effects of mutations elsewhere on the chromosome. Therefore, we complemented the *phoP* mutation using the pRMC2 vector (pRMC2:*phoP*). Expression of *phoP* in the *phoP*::tn mutant restored cytotoxicity to wild-type levels (Fig. 2a). As THP-1 cells are particularly sensitive to lysis by PSMs in the *S. aureus* supernatant, we hypothesised that PSM abundance may be reduced in the supernatant of the *phoP::tn* mutant. To test this, we extracted the bacterial supernatant and analysed PSM abundance using SDS-PAGE (Fig. 2b and Supplementary Fig. 1). An *agrA::tn* mutant, which does not produce any PSMs, was used as a control. We found that the *phoP::tn* mutant has a lower abundance of PSMs in the supernatant compared to the wild-type strain, explaining the reduced cytotoxicity of this strain. Additionally, this phenotype could also be complemented by expressing *phoP* from the pRMC2 vector.

**Figure 2.**
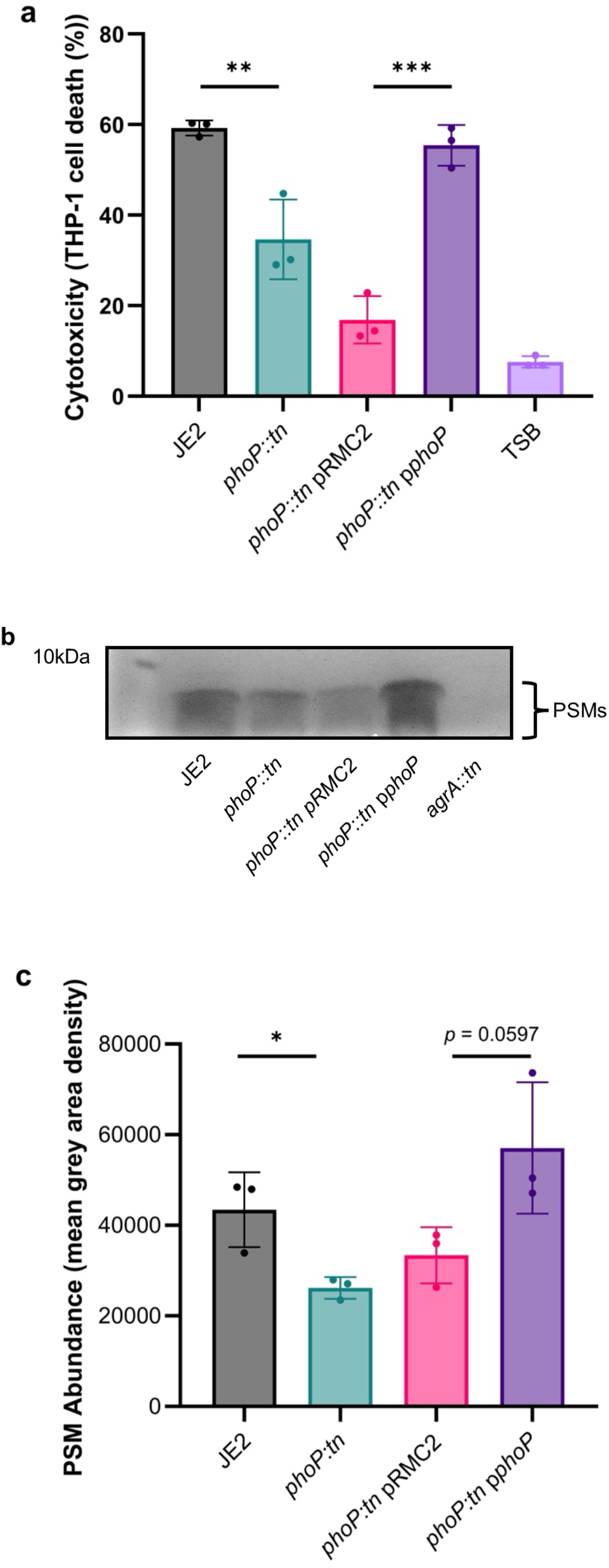
The *phoP::tn* mutant has a lower abundance of PSMs in the supernatant. (a) THP-1 cell lysis upon incubation with bacterial supernatant was quantified using trypan blue exclusion. The *phoP::tn* mutant exhibits lower cytotoxicity to THP-1 cells, an effect which can be complemented by expressing *phoP* from the pRMC2 plasmid. (b + c) Bacterial supernatants were extracted, separated via SDS-PAGE and protein bands visualised using Coomassie staining. (b) A representative SDS-PAGE gel showing PSMs in strains, which are visualised just below the 10kDa molecular weight marker, as shown previously (41-43). The band is absent in an *agrA::tn* mutant, which does not produce PSMs and a lower abundance can be seen in the *phoP::tn* mutant, which can be complemented. (c) Mean grey area of PSMs band calculated using densitometry. Significance in Fig. 2a + c is denoted as \**p* < 0.05, ** < 0.01, *** < 0.001.

### The *phoP::tn* Mutant has Reduced Agr Activity

Given that the Agr TCS is known to directly regulate expression of PSMs (22), we hypothesised that the activity of the Agr system was reduced in the *phoP::tn* mutant. To test this, we introduced a RNAIII::*gfp* fusion plasmid, which acts as a reporter of Agr activity, into the wild-type strain and *phoP::tn* mutant and measured fluorescence and growth over a 24h period. An *agrA::tn* mutant, which has no Agr activity, was used as a control. Despite minor differences in the growth curve, there was significant reduction in fluorescence in the *phoP::tn* mutant, demonstrating that RNAIII transcription was reduced in comparison to the wild-type strain. This explains the observed reduced cytotoxicity and PSM abundance in the *phoP::tn* mutant.

### PhoP Differentially Regulates Agr Activity Depending on Phosphate Availability

As PhoPR is activated by Pi limitation, we wanted to determine whether PhoP alters Agr activity in response to Pi availability. Induction of *pstS* expression, a phosphate-binding protein in the *pstSCAB* operon, can be used as a marker of PhoPR activation and Pi limitation as *pstS* expression is induced by phosphorylated PhoP (17, 44). To establish a model of Pi limitation, we introduced a *pstS*::*yfp* reporter fusion plasmid into the wild-type strain and *phoP::tn* mutant. We then compared fluorescence of the wild-type strain and *phoP*::tn mutant following an overnight growth in phosphate-depleted RPMI media supplemented with a range of Pi concentrations (Fig. 4a). In all Pi concentrations we tested (ranging from 0.125mM – 6mM), the wild-type and *phoP::tn* mutant grew to a similar OD_600nm_ (Supplementary Fig 2a + b). We detected *pstS* induction in the wild-type strain when Pi availability was ≤0.25mM. In contrast, *pstS* induction was absent in the *phoP::tn* mutant. Next, we measured Agr activity in the same growth conditions. When Pi availability was relatively high (between 1-6mM Pi), the *phoP::tn* mutant had significantly reduced Agr activity compared to the wild-type strain. This was similar to what was observed in TSB (Fig. 3b), which is a phosphate-rich media (∼14mM Pi). However, when Pi availability was ≤0.25mM we found that the *phoP::tn* mutant had significantly higher Agr activity compared to the wild-type strain. Furthermore, Agr activity increased in the WT strain between 0.125mM and 6mM (*p* = 0.0027), further supporting the notion that Pi availability regulates Agr activity. Overall, our data supports a model whereby PhoP exhibits differential regulation of Agr activity depending on Pi availability and its phosphorylation state. When Pi availability is high, PhoP is unphosphorylated and upregulates Agr activity. In contrast, when Pi availability is low and PhoP is phosphorylated, it represses Agr activity.

**Figure 3.**
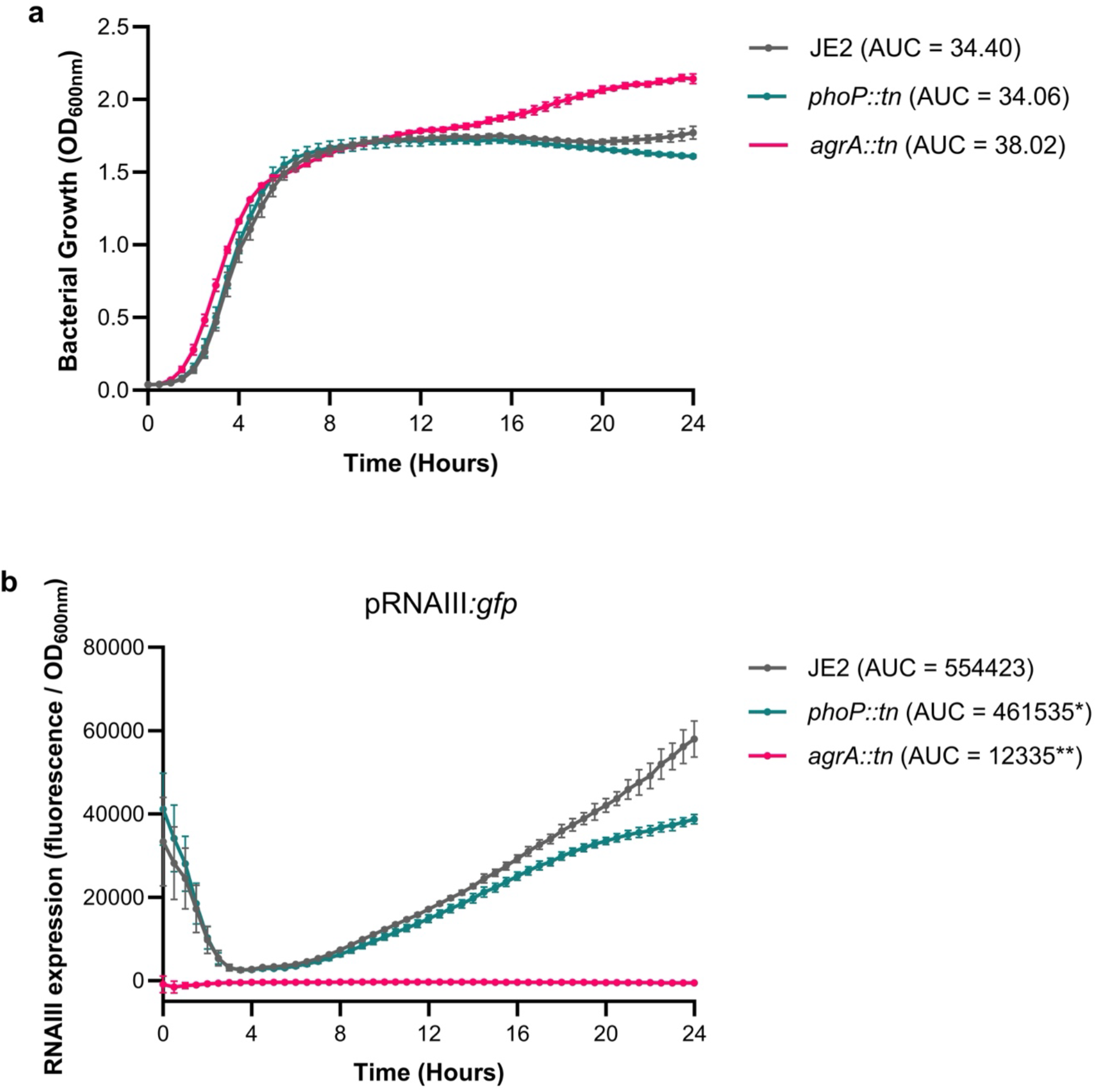
Activity of the Agr system is reduced in the *phoP::tn* mutant. (a) Growth of JE2 and the *phoP::tn* mutant carrying an RNAIII::*gfp* fusion plasmid was measured over 24 hours, demonstrating mutation of *phoP* has no effect on *S. aureus* growth. An *agrA::tn* mutant was used as a control. (b) Fluorescence (F) of JE2 and the *phoP::tn* mutant was measured to quantify activation of the Agr system. The *phoP::tn* mutant exhibited reduced fluorescence over 24 hours, indicating that this mutant has impaired Agr activity. Area under the curve (AUC) analysis was performed on OD600nm and F/OD values and significance determined using a *t*-test. Significance is denoted as \**p* < 0.01 and ** < 0.0001.

**Figure 4.**
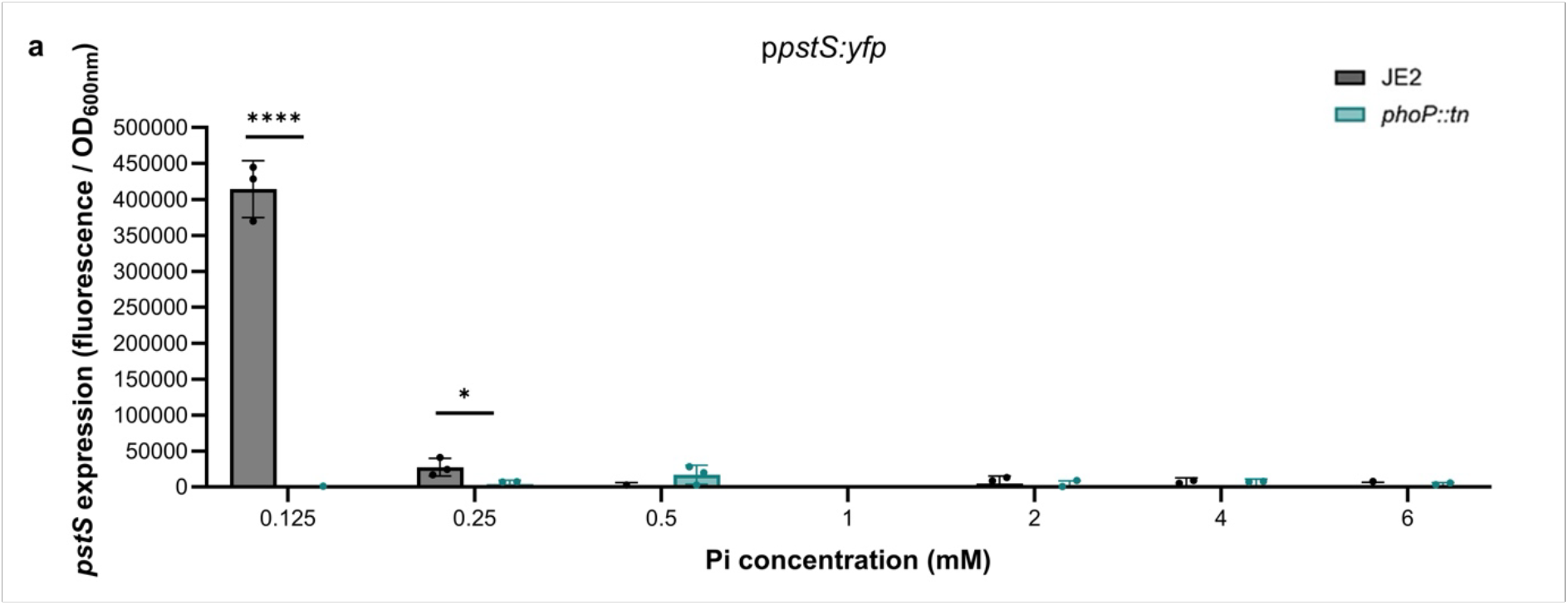
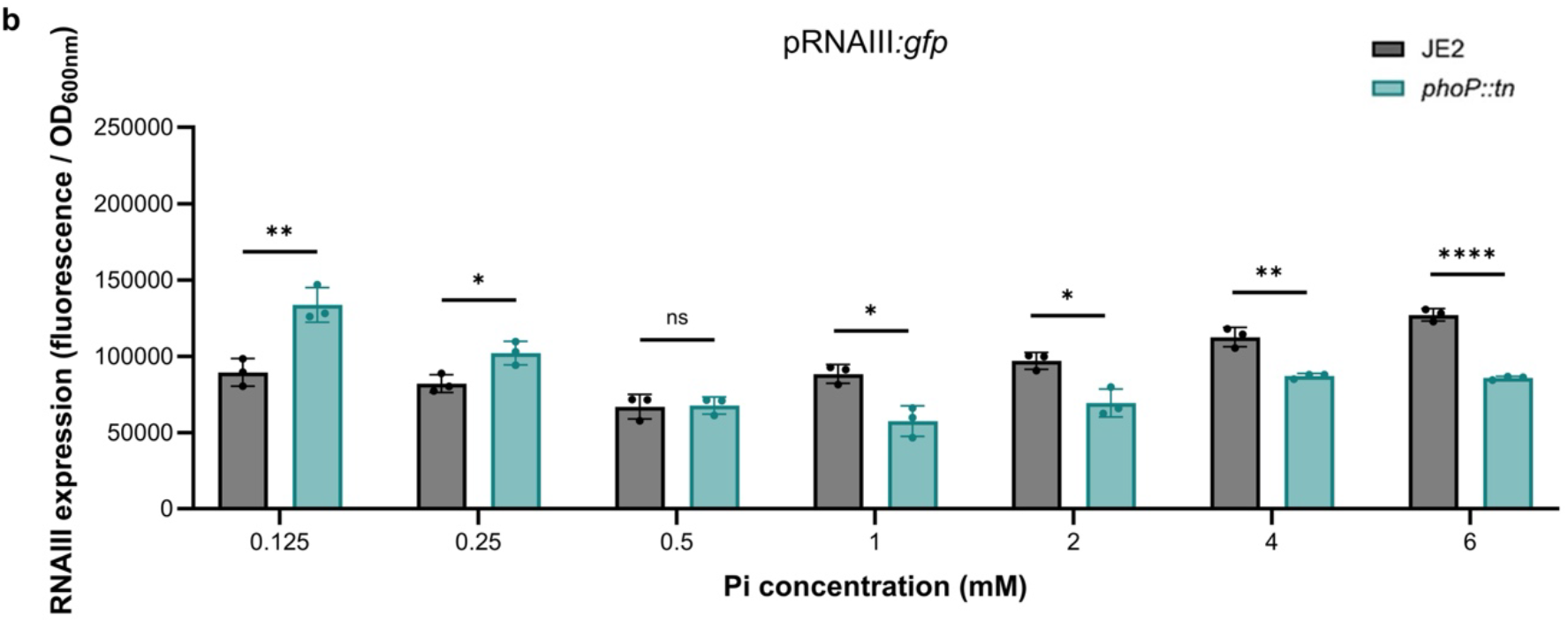
PhoP differentially regulates Agr Activity depending on phosphate availability. (a) Fluorescence (F) of JE2 WT and *phoP::tn* mutant carrying a *pstS*::*yfp* reporter plasmid was measured following growth in phosphate-depleted RPMI media supplemented with a range of inorganic phosphate (Pi) concentrations to identify PhoPR activation. Expression of *pstS* occurs below 0.25mM Pi and this induction does not occur in the *phoP::tn* mutant. (b) Fluorescence of JE2 and the *phoP::tn* mutant carrying an RNAIII::*gfp* reporter plasmid was measured in the same condition to determine Agr activity. The *phoP::tn* mutant had reduced fluorescence at high Pi concentrations, but higher fluorescence Pi limitation compared to the wild-type strain. Each dot represents one biological replicate (n = 3), error bars the standard deviation, and statistical significance was determined using a *t* test. Significance is denoted as \**p* < 0.05; ** < 0.01; < ***0.001.

### PhoP represses cytotoxicity in low Pi environments

Our data demonstrates that PhoP acts as an activator in high Pi environments as the *phoP::tn* mutant has lower Agr activity and cytotoxicity and overexpression of *phoP* increases cytotoxicity in these growth conditions. In contrast, we found that the *phoP::tn* mutant has higher Agr activity in low Pi environments (≤0.25mM), suggesting it may act as a repressor in this environment. To confirm this, we tested the cytotoxicity of strains following growth in either 0.125mM or 0.25mM Pi (Fig. 5a + b). In line with Agr activity under low Pi conditions, the *phoP::tn* mutant had significantly higher cytotoxicity compared to the wild-type strain in both 0.125mM or 0.25mM Pi. Furthermore, expression of *phoP* in the *phoP*::tn mutant significantly repressed cytotoxicity, confirming that PhoP acts as a cytotoxicity repressor in low Pi environments.

**Figure 5.**
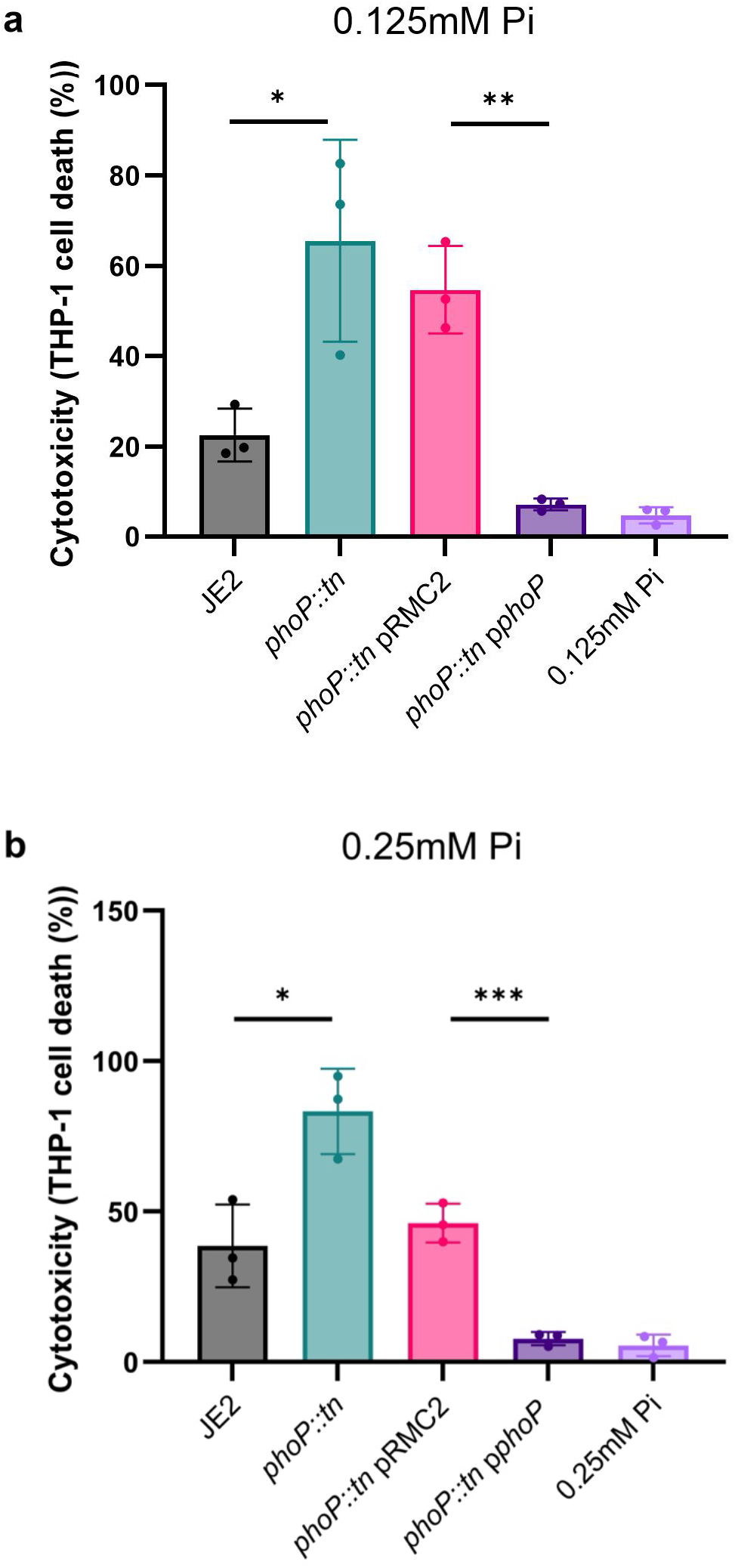
PhoP acts as a repressor of cytotoxicity in low Pi environments. (a + b) THP-1 cell lysis upon incubation with bacterial supernatant was quantified using trypan blue exclusion. Supernatants were extracted following growth of strains in phosphate-depleted RPMI media supplemented with either 0.125mM Pi (a) or 0.25mM Pi (b). In both conditions, the *phoP::tn* mutant has higher cytotoxicity compared to the wild-type strain and expression of phoP from the pRMC2 significantly represses cytotoxicity compared to the *phoP::tn* mutant complemented with an empty pRMC2 vector. Significance is denoted as \**p* < 0.05; ** < 0.01; < ***0.001

### Regulation of Agr activity by PhoP is Determined by its Phosphorylation State

As we identified PhoP can act as either an activator or repressor of cytotoxicity depending on Pi availability, we wanted to verify the impact of PhoP phosphorylation on Agr activity. To test this, we substituted the phosphorylation site of PhoP (D53) in pRMC2::*phoP* with an alanine, which prevents phosphorylation (phosphoablative), or a glutamic acid residue, which mimics the conformational change induced by permanent phosphorylation of the PhoP protein (phosphomimetic) (17). We then induced expression of either D53A or D53E PhoP and measured expression of *pstS* and RNAIII. As the pRMC2 and pRNAIII::*gfp* vectors use the same resistance marker, we measured the levels of *pstS* and RNAIII using qRT-PCR.

Transcription of *pstS* was elevated in *phoP:tn* expressing either D53A or D53E PhoP compared to *phoP:tn* pRMC2. However, this was 1146-fold higher upon expression of D53E PhoP compared to D53A PhoP, suggesting that induction of *pstS* expression is mediated by phosphorylated PhoP. Transcription of RNAIII was also elevated in both strains. However, expression of D53A PhoP resulted in approximately 2-fold higher RNAIII transcription compared to D53E PhoP. Combined, this data confirms that phosphorylation state affects PhoP regulation of Agr activity.

**Figure 6.**
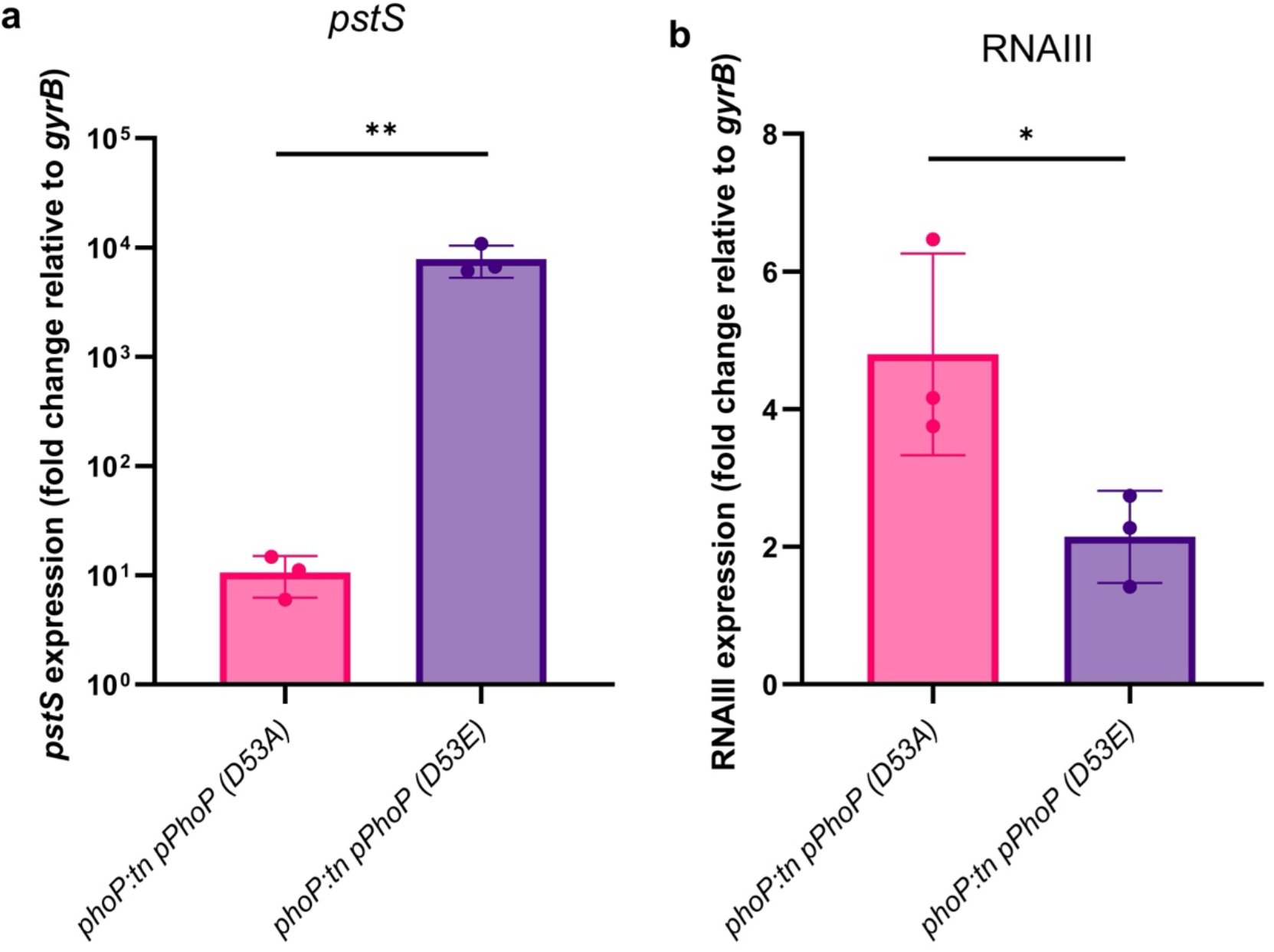
Phosphorylation state of PhoP determines Agr regulation. Gene expression of either *pstS* (a) or RNAIII (b) was quantified relative to expression levels of the *gyrB* housekeeping gene in strains expressing D53A or D53E PhoP. Fold change was then calculated relative to the expression of each target gene in a strain carrying the empty pRMC2 vector. Expression of *pstS* is significantly higher when expressing D53E PhoP compared to D53A PhoP. In contrast, expression of D53A PhoP resulted in higher levels of RNAIII expression compared to expression of D53E PhoP. Each dot represents one biological replicate (n = 3), error bars the standard deviation, and statistical significance was determined by a *t*-test. Significance is denoted as \**p* < 0.05 and \*\**p* < 0.01.

## Discussion

In this study, we systematically compared the contribution to cytotoxicity of all the non-essential TCSs in the genome of *S. aureus* strain JE2. Our screening identified a link between TCSs involved in cell wall biosynthesis or autolysis and regulation of cytotoxicity as several mutants (*graR::tn, graS::tn, nsaR::tn* and *lytR::tn*) had reduced THP-1 toxicity. Disruption of these TCSs has profound effects on cell surface properties and morphology (18, 45-48). Accordingly, changes to the cell surface structure of *S. aureus* have been previously reported to affect the regulation and secretion of toxins (35, 49). Nutrient sensing was also identified as an important signal for regulating cytotoxicity with *kdpE::tn, phoP::tn* and *hptS::tn* mutant exhibiting reduced cytotoxicity. As the biosynthesis of cytolytic toxins is an energetically costly process, many pathogenic bacteria synchronise this with nutrient availability (14, 50). Furthermore, nutrient availability can vary between different host tissues, acting as a marker for host environments where cytotoxicity is an advantageous phenotype.

Having decided to focus here on the effect of PhoPR on cytotoxicity, we demonstrated that the *phoP::tn* mutant produces less PSMs in the bacterial supernatant during growth in TSB. PSMs include PSMα1-4 and PSMβ1/2, which are directly regulated by phosphorylated AgrA. Additionally, delta-haemolysin (Hld), is encoded within the main effector molecule of the Agr system, RNAIII (51, 52). We identified that the *phoP::tn* mutant has lower RNAIII expression, suggesting that this mutant has reduced amounts of Hld and lower Agr activity. This finding supports the notion that TCSs in *S. aureus* are a highly interconnected network which frequently interact with each other to fine-tune gene expression (3, 53). For example, expression of a phosphomimetic form of WalR activates the Sae TCS (54). Similarly, glucose-6-phosphate induces expression of both Agr and Sae and this is dependent on the presence of the HptRS TCS (55).

Induction of *pstS* by PhoP follows the canonical signalling mechanism by which a two-component system activates a target gene. For example, we detected induction of *pstS* expression in growth media ≤0.25mM Pi, where PhoPR is activated and levels of phosphorylated PhoP significantly increase (17, 44). In contrast, no regulation of *pstS* was detected when Pi levels were ≥0.5mM, where PhoP is unphosphorylated. This mechanism was confirmed through expression of D53E PhoP, which resulted in 1146-fold higher levels of *pstS* compared to expression of D53A PhoP. However, PhoP-mediated regulation of Agr activity follows an atypical trend. Firstly, unphosphorylated PhoP acts as a strong activator of RNAIII transcription in high Pi environments. This allows for regulation of Agr activity in the absence of an activation signal and explains why a *phoR::tn* mutant did not show a reduction in cytotoxicity in our initial screening as the experiment was performed at high Pi. Secondly, we identified that phosphorylated PhoP has differential effects on Agr activity depending on Pi concentration, acting as a weak activator in high Pi and a repressor in low Pi. Our current hypothesis is that additional factors expressed at low Pi regulated by PhoP concur in the regulation of Agr activity. This would explain why Agr activity is elevated in a *phoP::tn* under Pi limitation and overexpression of PhoP in this condition strongly represses cytotoxicity. This also suggests that regulation of cytotoxicity by Pi is a multifactorial process, with additional factors involved in coordination with PhoPR.

**Figure 7.**
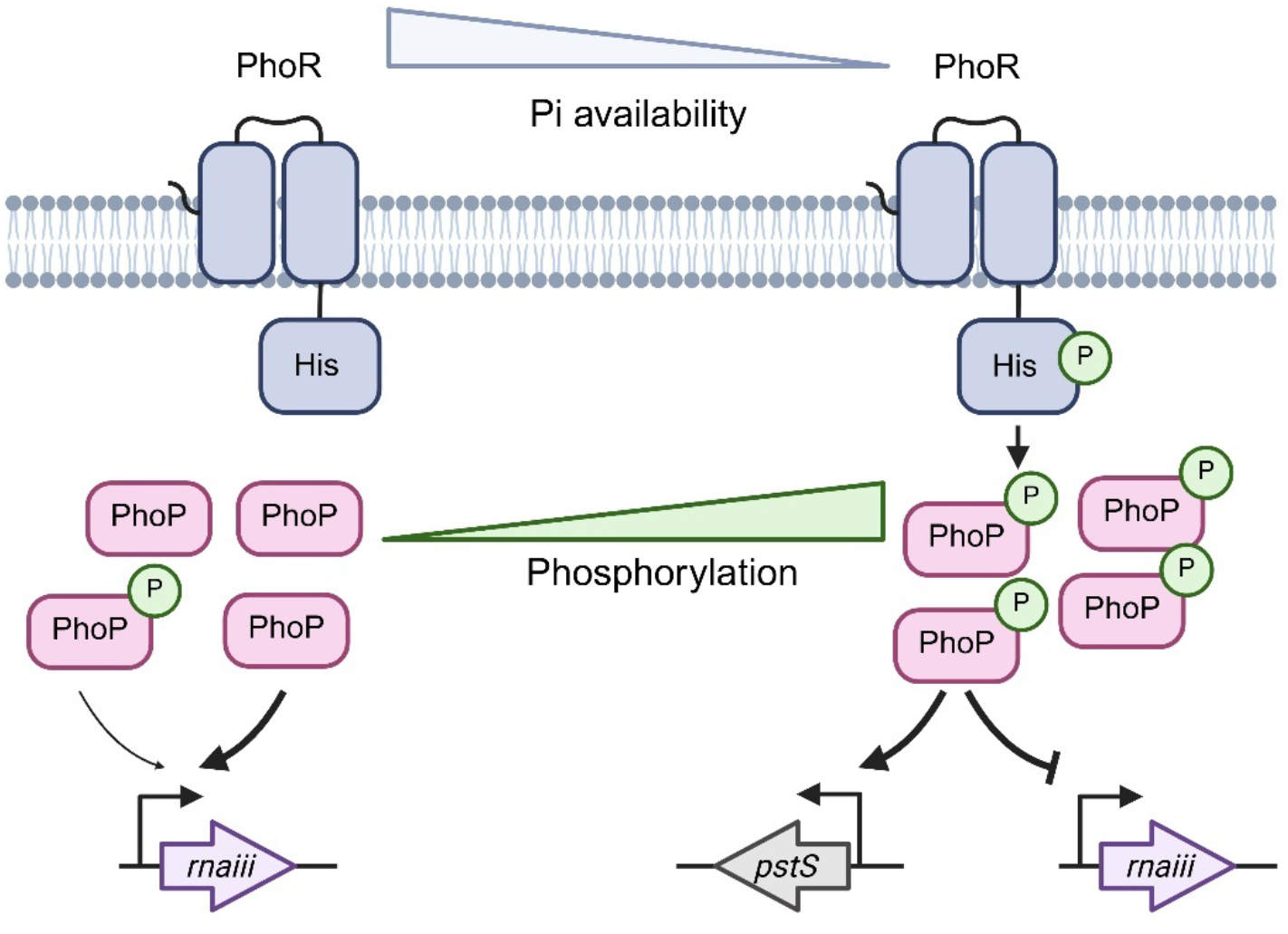
Summary of Agr regulation by PhoP. When phosphate (Pi) availability is high, PhoR is inactive and PhoP mainly exists in its unphosphorylated form. Unphosphorylated PhoP acts as a strong activator of Agr activity. Phosphorylated PhoP also acts as a weaker activator of Agr activity. When Pi availability decreases, activation of PhoR occurs. PhoR undergoes autophosphorylation and mediates phosphorylation of PhoP. Phosphorylated PhoP induces expression of genes involved in phosphate transport, including *pstS*. Furthermore, phosphorylated PhoP acts as a repressor of Agr activity in low Pi environments.

The finding that unphosphorylated PhoP is a stronger activator of RNAIII than phosphorylated PhoP in high Pi environments adds to growing experimental evidence which indicates unphosphorylated RRs have important regulatory functions (56). For example, unphosphorylated WalR from *Streptococcus pneumoniae* inhibits *fabT*, a repressor of fatty acid chain elongation (57). This supports the idea that TCSs can initiate a more complex response than a simple on-off feed forward mechanism, providing several regulatory options depending on their phosphorylation status. One more potential layer of regulation could be represented by the expression levels of PhoP, which may also change depending on Pi availability and affect how PhoPR regulates Agr activity.

To date, studies of PhoPR in *S. aureus* have focused on its role in regulating phosphate transporters (*pstSCAB, nptA* and *pitA*) in response to phosphate limitation (44). Importantly, a *S. aureus* Δ*phoPR* mutant was found to be attenuated in the heart during a systemic staphylococcal abscess model of infection whereas a Δ*pstSCAB* Δ*nptA* double mutant had no defect, suggesting transporter-independent factors contribute to *S. aureus* pathogenesis in this environment (44). Our findings that PhoPR regulates cytotoxicity provides a potential explanation for this virulence defect and suggests that this regulation may be relevant during *in vivo* infections.

In conclusion, our systematic screening approach revealed how the regulatory effects of 11 TCS out of 16 converge on cytotoxicity. Our results confirm that virulence regulation depends on an intricate regulatory network that reacts to diverse environmental conditions. We identified phosphate sensing via PhoP as a novel regulator of cytotoxicity in *S. aureus*. PhoP interacts with the Agr system to regulate expression of PSMs. Crucially, PhoP mediates differential regulation of RNAIII expression depending on environmental Pi and the balance between its unphosphorylated and phosphorylated forms. This work builds upon previous studies which have identified the PhoPR TCS as important for the full virulence of *S. aureus* and inhibition of this system represents a promising therapeutic target.

