## Supplementary table and data for "Phosphate sensing by PhoPR regulates the cytotoxicity of *Staphylococcus aureus*"

**Supplementary Figures**

| Strain | *p*-value | Adjusted *p*-value |
|---|---|---|
| *agrA* | 0.000004 | 0.000041 |
| *agrC* | <0.000001 | 0.000009 |
| *saeR* | 0.000391 | 0.001579 |
| *saeS* | 0.000805 | 0.002324 |
| *arlR* | 0.004718 | 0.007340 |
| *arlS* | 0.000292 | 0.001475 |
| *graR* | 0.009111 | 0.010826 |
| *graS* | 0.006006 | 0.008666 |
| *vraR* | 0.038405 | 0.043099 |
| *vraS* | 0.106537 | 0.097820 |
| *nsaR* | 0.000769 | 0.002324 |
| *nsaS* | 0.000077 | 0.000518 |
| *lytR* | 0.003554 | 0.007180 |
| *lytS* | 0.061038 | 0.064893 |
| *kdpE* | 0.004303 | 0.007340 |
| *kdpD* | 0.191049 | 0.160800 |
| *phoP* | 0.001853 | 0.004423 |
| *phoR* | 0.077734 | 0.078511 |
| *hptR* | 0.227762 | 0.184032 |
| *hptS* | 0.001971 | 0.004423 |
| *hssR* | 0.124835 | 0.109638 |
| *hssS* | 0.411666 | 0.307987 |
| *srrA* | 0.639063 | 0.461038 |
| *srrB* | 0.783256 | 0.545578 |
| *nreC* | 0.004723 | 0.007340 |
| *nreB* | 0.381898 | 0.296706 |
| *airS* | 0.101433 | 0.097569 |
| *desR* | 0.007443 | 0.009396 |
| *desK* | 0.006940 | 0.009346 |

Supplementary Table 1. Statistical analysis of cytolytic activity in TCS transposon mutants. Significance was determined using multiple t-tests with a false discovery rate (FDR) of 1% applied. Differences in cytotoxicity were considered significant if they had an adjusted p-value ≤0.01.

**a**

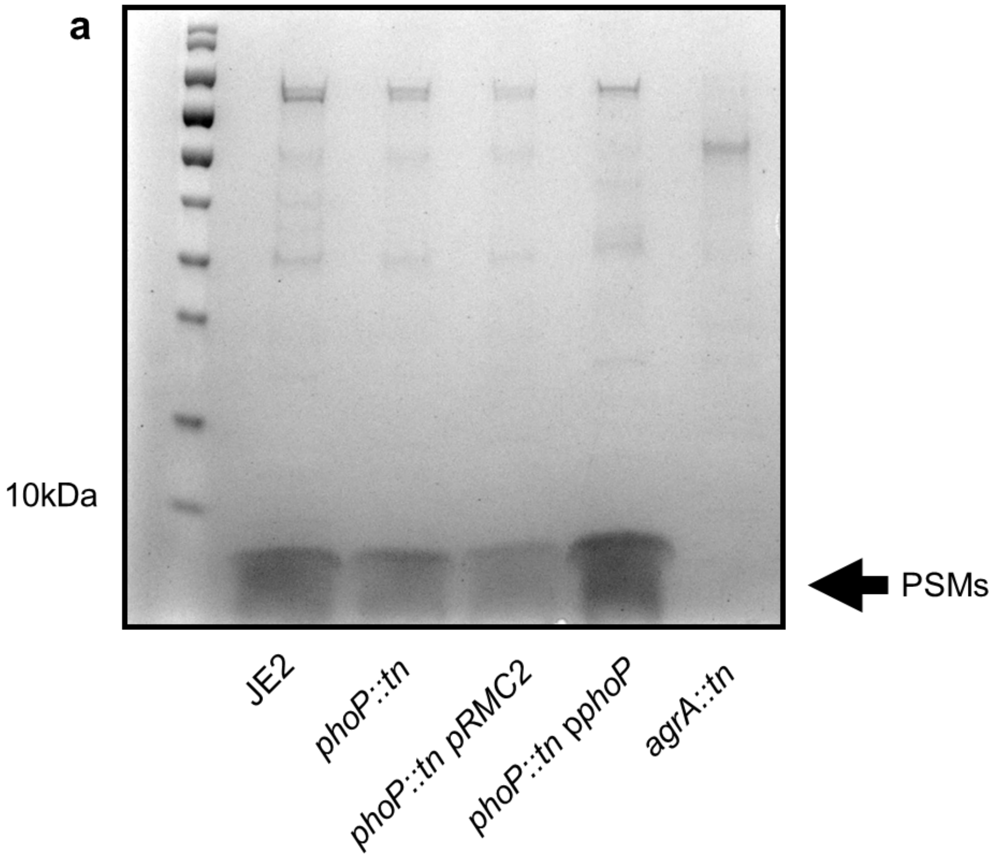
10kDa
PSMs
JE2
phoP::tn
phoP::tn pRMC2
phoP::tn pphoP
agrA::tn

**b**

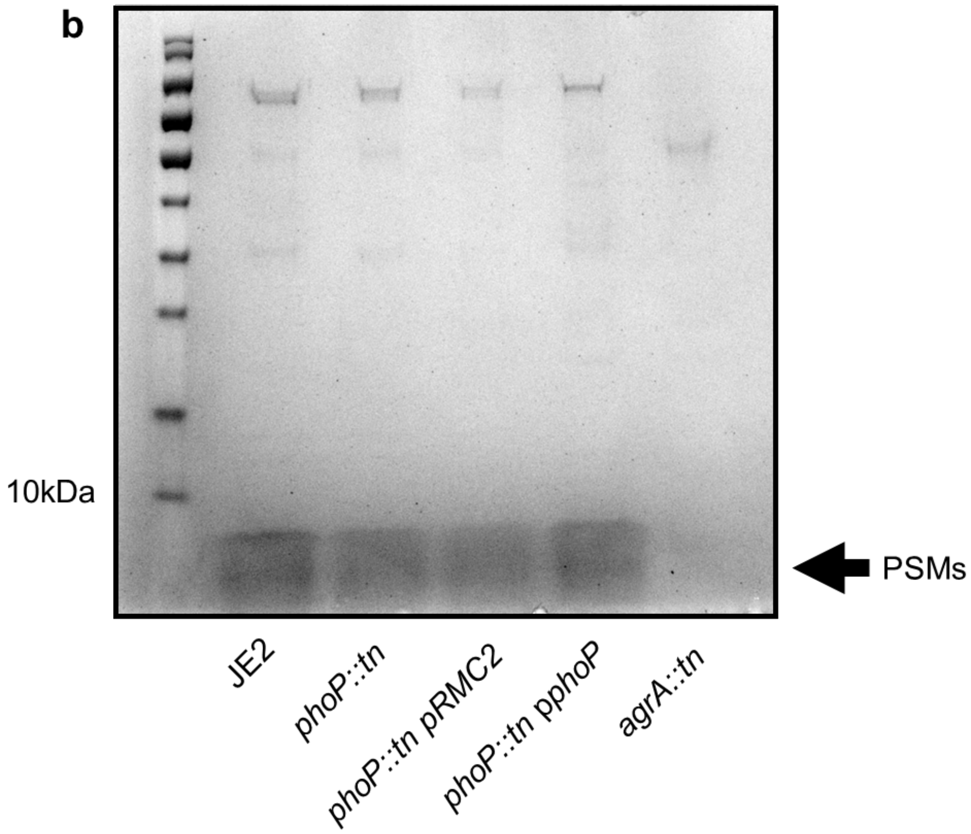
10kDa
PSMs
JE2
phoP::tn
phoP::tn pRMC2
phoP::tn pphoP
agrA::tn

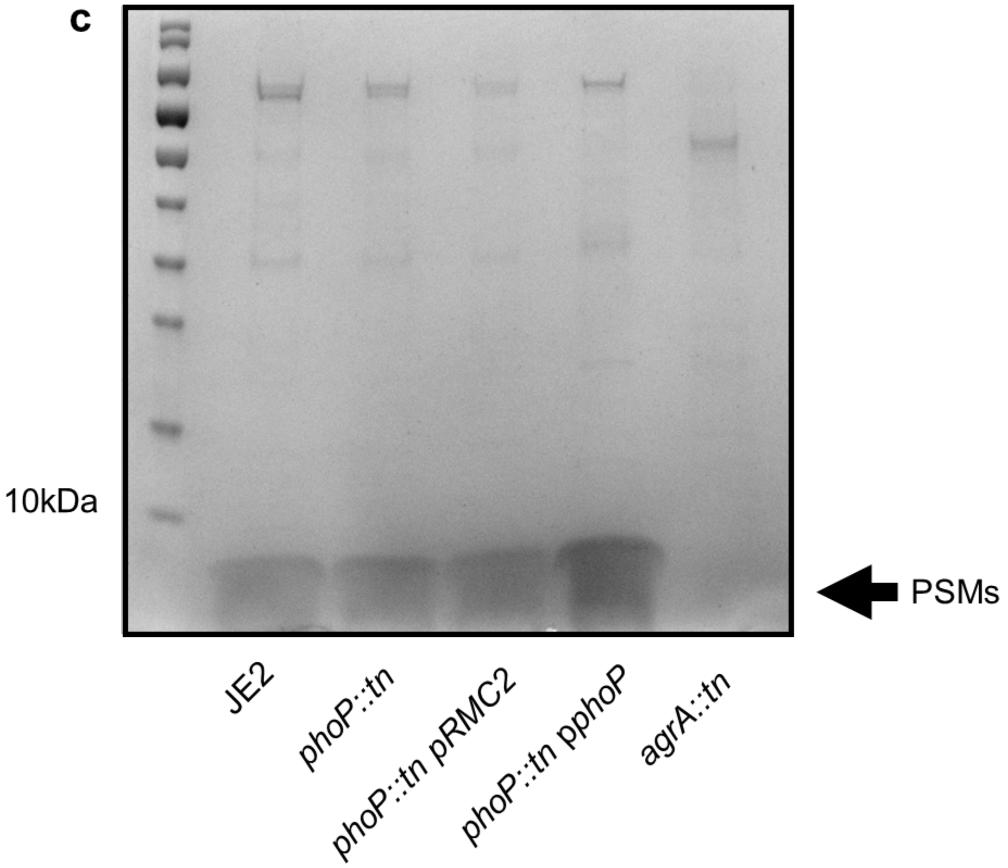


Supplementary Figure 1. Three replicates (a,b and c) of SDS-PAGE gels showing a reduction in PSMs in the supernatant of the *phoP::tn* mutant, which can be complemented by reintroducing expression of *phoP*.

**a**

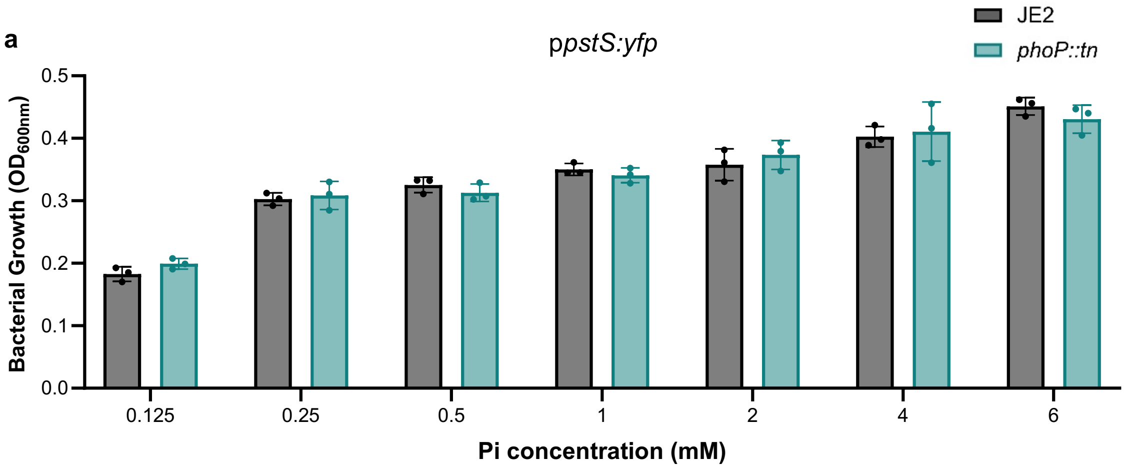


**b**

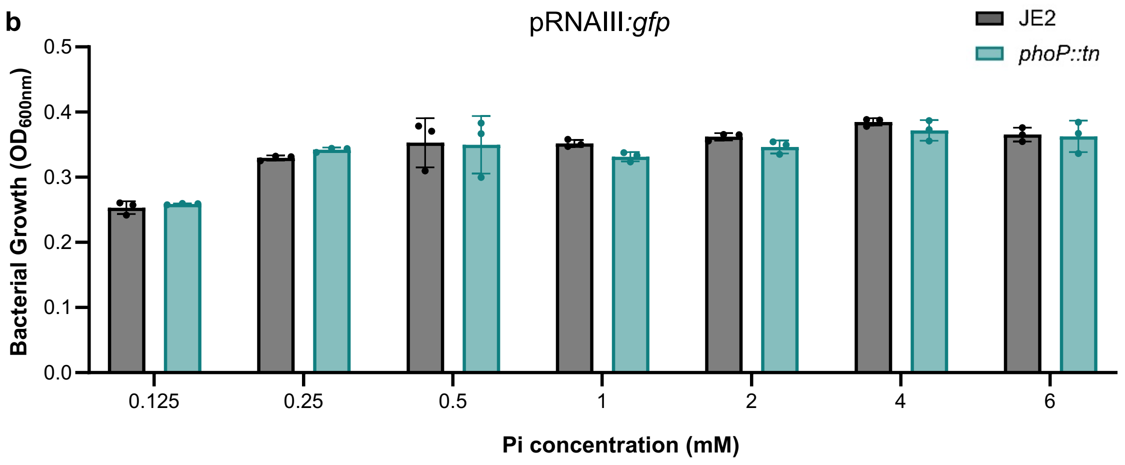


Supplementary Figure 2. Growth of JE2, the *phoP::tn* mutant carrying *ppstS::yfp* (a) or p*rnaiii::gfp* (b) was measured from an overnight culture in RPMI media with different phosphate concentrations. All three strains grew to a similar $OD_{600nm}$ in all concentrations tested.
